# Characterization of a chronic UV-induced photoaging mouse model: insights into skin barrier dysfunction, extracellular matrix remodeling, and altered adipogenesis

**DOI:** 10.64898/2026.04.10.712660

**Authors:** Martina Bajerová, Romana Šínová, Matěj Šimek, Kateřina Lehká, Petra Ovesná, Martin Čepa, Iva Dolečková, Vladimír Velebný, Kristina Nešporová, Lukáš Kubala

**Affiliations:** Institute of Biophysics of the Czech Academy of Sciences, Brno, Czech Republic; Contipro a.s., Dolní Dobrouč 401, 56102, Dolní Dobrouč, Czech Republic; Institute of Biostatistics and Analyses, Masaryk University, Kamenice 5, Brno, 625 00, Czech Republic; Institute of Experimental Biology, Faculty of Science, Masaryk University, Brno, Czech Republic; International Clinical Research Center, St. Anne’s University Hospital Brno, Brno, Czech Republic

**Keywords:** hairless mice, photoaging, inflammation, hyaluronan, adipogenesis, TSG-6

## Abstract

Chronic exposure to ultraviolet (UV) radiation, known as photoaging, accelerates skin aging by inducing molecular, histological, and functional changes. This study established a mouse model using SKH-1 hairless mice to investigate chronic UV-induced photoaging over eight weeks. SKH-1 hairless mice were exposed to a combination of UVA and UVB, and the progression of skin damage was monitored through physical, histological, and molecular parameters, with a focus on erythema, transepidermal water loss, and collagen and hyaluronan (HA) metabolism. Significant reductions in HA content and alterations in DNA repair markers, such as γH2AX, were observed, highlighting the impact of chronic UV exposure on skin structure and function. Reactive adipogenesis and increased epidermal thickness were noted, reflecting adaptive responses to UV-induced damage. By investigating these parameters over the evaluation period, we provide a comprehensive time-course analysis of the progressive impact of UV-induced photoaging, offering insights into the underlying mechanisms and potential therapeutic targets to prevent or delay photoaging.

## 1. Introduction

UV radiation-induced skin aging, or photoaging, is one of the main external factors that mediate pathological changes in the skin related to age. It might account for up to 90% of visible skin changes by inducing molecular and cellular changes that result in wrinkles, discoloration, and sagging skin.^1^ Photoaging is directly correlated with an increased risk of cancer due to UV radiation-induced impairment of DNA repair enzymes, immune functions, and other factors.^2^ Understanding the molecular and cellular processes occurring during photoaging in the complex context of skin tissue, which consists of many cell types in a highly intricate, multilayered arrangement, is crucial for any future research on this pathological phenomenon.

UV radiation wavelengths relevant to regular sunlight have different effects mediated by different mechanisms. UVA radiation penetrates the dermis, causing structural changes linked to aging. While most UVB light is absorbed by the skin’s outermost layer, the remaining 30% reaches deeper layers, making it a primary cause of skin erythema and sunburn, which significantly contribute to photoaging and increase the risk of skin cancer development.^3^ Increased reactive oxygen species production triggers pro-inflammatory and degradative reactions, disrupting the collagen structure and blocking procollagen synthesis.^4^ Pro-inflammatory cytokines such as interleukin-1 (IL-1), IL-6, IL-8, and tumor necrosis factor α (TNF-α) and matrix metalloproteinases (MMPs), which hydrolyze collagen fibrils, are involved in these processes.^2,4,5^ Thus, among the major effects of chronic UV radiation are alterations in key components of the extracellular matrix (ECM), particularly collagen and hyaluronic acid (HA). HA is a glycosaminoglycan that is essential for maintaining skin hydration, elasticity, and overall integrity. Collagen fibers break and form dispersed clusters, leading to the loss of ECM integrity in photoaged skin. Chronic UV exposure depletes HA in the skin, reducing moisture retention and compromising the skin barrier. The breakdown of these ECM components, along with other components such as elastin, accelerates the formation of wrinkles and the skin sagging, further contributing to the aging phenotype.^6^ Recent studies have reported that UV radiation disrupts lipid metabolism in the deeper layers of skin and induces alterations in adipogenesis.^7,8,9,10^

To study photoaging, laboratory mice can be used. Typically, photoaging is induced by exposing mice to repeated UV radiation for several weeks.^11^ SKH-1 hairless mice are immunocompetent and hair-free, making them ideal for UV radiation experiments.^12,13^ Other strains, such as C57BL/6 or BALB/c, are also used, but they might require shaving, risking skin irritation. Despite the extensive use of mice in photoaging research, significant inconsistencies persist owing to variations in experimental design, including the duration, UV dose, wavelength, frequency, and differences in the sensitivity of mouse strains to UV. Additionally, many studies analyze parameters at only a single time point, limiting the ability to compare molecular mechanisms or tested compounds across different studies. These discrepancies highlight the need for a standardized, well-characterized model, which we address in this study by examining multiple time points and comprehensive endpoints.

The aim of this work was to introduce and thoroughly characterize an animal model of skin aging caused by chronic UV irradiation that would be suitable for robust exploration of photoaging mechanisms and pharmacological interventions. Using SKH-1 mice, we aimed to characterize the physical, histological, and molecular changes occurring in the irradiated skin during the 8-week period. A specific focus was on skin barrier function, collagen content, histological changes, and adipogenesis. Additionally, we evaluated DNA damage, the levels of pro-inflammatory eicosanoids, and the expression of genes involved in inflammation and tissue repair processes.

## 2. Materials and methods

### 2.1. Animals and animal care

All the animal studies were conducted in compliance with ethical guidelines under the approval of the Czech Academy of Sciences (CAS) (file number: AVCR-S 1114/2022 SOV II, 01 July 2022) and were supervised by the local ethical committee of the Institute of Biophysics of the CAS.

SKH-1 hairless mice were obtained from Velaz s.r.o. (Czech Republic). The mice were bred and housed in individually ventilated cages with free access to sterile drinking water and standard food in a certified animal facility at the Institute of Biophysics of the CAS (Czech Republic), with a controlled climate, hygiene, humidity, temperature, and a 12-hour light/dark cycle. Eight-week-old female mice were used in the experiment due to their lower aggression levels when co-housed, minimizing the risk of scratches or bites, which could interfere with the skin integrity. Mice were assigned to eight groups using stratified randomization based on the initial weights (16-26 g) to ensure comparable weight distribution across groups. Four groups served as nonirradiated controls and were sacrificed at 2 (N=6), 4 (N=6), 6 (N=6), and 8 (N=5) weeks. The remaining four groups were irradiated (described below) and sacrificed at the same intervals: 2 (N=6), 4 (N=6), 6 (N=6), and 8 (N=7) weeks post-exposure.

### 2.2. UV irradiation

UV irradiation was provided by a 300 W xenon arc lamp (Oriel/Newport, with FSQ-UG5 and 20CGA-280 filters to simulate solar UV at 280–410 nm). The UVB intensity at a depth of 14.5 cm was measured with a PMA2100V radiometer. The intensity of UVB radiation, measured at a distance of 14.5 cm from the irradiated surface, was measured via a Photometer/Radiometer PMA2100V detector (Solar Light Company, Inc.). Mice were anesthetized by inhalation of isoflurane (2.5–5.0%) during each irradiation session to minimize movement and ensure standardized exposure. A custom-made cardboard mask with a 2×2 cm square window was placed over each animal, exposing a defined area of the dorsal skin to UV light while shielding the rest of the body. To ensure uniformity of irradiance, each mouse was individually removed from its cage and positioned under the UV lamp. This individualized setup, combined with controlled anesthesia and masking, ensured reproducible and consistent UV exposure across all treatment groups.

Before the experiment was initiated, the minimal erythema dose (MED) was estimated by testing four different UV doses (25, 50, 75, and 100 mJ/cm²) on the backs of two SKH-1 mice, with two sites tested per mouse. Each test area as well as control nonirradiated area on each mouse was inspected, weighted (Supplementary Figure 1), photographed, and measured at 24, 48, and 72 hours post-irradiation.

The dorsal skin of each mouse was irradiated three times a week for a total duration of 2, 4, 6, or 8 weeks. The irradiance began at 1 MED and gradually increased to 4 MED over the first 4 weeks. This level of irradiation (4 MED) was maintained for the rest of the experiment. The gradual increase in MED was inspired by the protocols described in previous studies.^14–16^ At the end of each exposure period, the mice were anesthetized by inhalation of isoflurane (2.5–5.0%). Blood samples were collected via cardiac puncture into 0.5M EDTA, serving as an anticoagulant. Skin samples were harvested and immediately immersed in RNA later for gene expression analysis, or fixed in 4% paraformaldehyde, or frozen in liquid nitrogen and stored at −80°C for further analysis.

### 2.3. Evaluation of erythema, skin hydration, and transepidermal water loss

To assess the effect of UV radiation on the skin, the MultiProbe Adapter System MPA 580 (Courage + Khazaka electronic GmbH, Germany) was used. The parameters were measured in the middle of the irradiated area and the corresponding area in non-irradiated control mice once a week before irradiation. The erythema index was determined via the Mexameter® MX18 probe (mean of at least 5 readings). The hydration of the *stratum corneum* (SC) was measured via a Corneometer® CM825 probe (mean of at least 5 readings). Transepidermal water loss (TEWL) was measured by the Tewameter® TM 300 probe (mean of at least 15 readings taken after saturation of the Tewameter chamber). Mice were anesthetized during the measurements to reduce the variability caused by movement. All these baseline skin parameters (erythema, TEWL, hydration) and body weight were measured prior to the first irradiation to confirm inter-group comparability.

### 2.4. Histological evaluation of the skin

Skin samples were fixed in 4% paraformaldehyde and histologically processed as described previously.^17^ The sections from the paraffin blocks were subjected to hematoxylinceosin (H&E) staining to examine epidermal and dermal thickness. To examine the abundance and density of collagen fibers, Masson’s trichome staining (Carl Roth, R.3459.1) was performed. The infiltration of neutrophils was analyzed by Naphthol AS-D chloroacetate esterase staining (Sigma Aldrich, 91C-1KT). A prepared solution consisting of nitride, Fast Red Violet LB Base, distilled water, TRIZMAL 6.3 buffer, and naphthol AS-D chloroacetate was applied to the deparaffinized slides for 45 minutes.

All samples were analyzed under a Leica DMi8 microscope with an objective HC PL FLUOTAR, 10x/0.32 DRY. Epidermal thickness was measured at five central locations of the sample and averaged. The number of positive purple cells (Naphthol AS-D chloroacetate esterase staining) was quantified via ImageJ.^18^ The threshold was set according to the purple signal and cell size. The green signal of collagen (Masson-trichrome staining) was quantified via QuPath 0.4.3.^19^ using color deconvolution and mean intensity quantification in ImageJ. Image analysis of the size and number of adipocytes was performed on H&E-stained skin sections. First, brightfield images were manually segmented to roughly separate the areas containing adipocytes. The segmented image was then transformed to grayscale, increased to the power of three, and multiplied by two. Since adipocytes appear brighter in the image, these arithmetic operations increase their brightness while simultaneously dimming the rest of the image, giving it an almost fluorescent-like appearance. Bright spots were then counted, and their size and form factor (the ratio of the circumference to the area normalized to a circle)^20^ were used to identify adipocytes and filter out similar-looking tissue regions. In addition to the initial manual segmentation, all the images were processed and analyzed via CellProfiler.^21^

### 2.5. Immunofluorescence analysis

Briefly, antigen retrieval (with Tris-EDTA buffer, pH 9, 95° C, 20 min) was performed on the skin slices, which were then permeabilized with 0.1% Triton X-100. The slides were incubated with blocking buffer (TBS with 10% fetal bovine serum, 0.1% Tween20, and 2.3% glycine) for 60 min at RT. The sections were subsequently incubated overnight at 4 °C with the following rabbit monoclonal primary antibodies: anti-phospho-histone H2A. X (1:200; Cell Signaling Technology, 9718), followed by Alexa Fluor 488-conjugated goat anti-rabbit IgG (1:500; Abcam, ab150077). The slices were mounted in ROTI Mount FluorCare DAPI (Carl Roth, HP20.1). Images were acquired via a Leica DMi8 fluorescence microscope with an objective HC PL FLUOTAR, 20x/0.40 DRY. Images of γH2AX were analyzed via Cellprofiler.^21^ γH2AX-positive cells were quantified as the number of DAPI-stained nuclei with colocalized green fluorescence signal corresponding to γH2AX, expressed as a percentage of the total number of nuclei.

### 2.6. Quantification of hyaluronic acid

HA was quantified via a validated liquid chromatographycmass spectrometry method as described previously.^17,22,23^

### 2.7. Quantification of eicosanoids

The quantification of selected eicosanoids, including 6-keto-prostaglandin F□α (6-keto-PGF□α), thromboxane 2 (TXB□), prostaglandin F□α (PGF□α), prostaglandin E□ (PGE□), prostaglandin D□ (PGD□), 5-hydroxyeicosatetraenoic acid (5-HETE), 12-hydroxyeicosatetraenoic acid (12-HETE), and 15-hydroxyeicosatetraenoic acid (15-HETE), was carried out via validated liquid chromatographycmass spectrometry methods as described previously.^17,22,23^

### 2.8. Gene expression analysis

The RNA from the skin samples was reversely transcribed to cDNA as described previously.^24^ Gene expression was measured via qPCR via the following TaqMan gene expression assays (Thermo Fisher Scientific): IL-1β Mm00434228_m1, MMP9 Mm00442991_m1, COL1A1 Mm00801666_g1, PPARγ Mm00440940_m1, adiponectin Mm04933656_m1, leptin Mm00434759, HAS2 Mm00515089_m1, HAS3 Mm00515092_m1, CEMIP Mm00472921_m1, CEMIP2 Mm00459599_m1, HYAL1 Mm00480053_m1, HYAL2 Mm00477731_m1, CD44 Mm01277161_m1, HMMR Mm00469183_m1, TNFAIP6 Mm00493736_m1 and Ribosomal Protein L13a - RPL13A Mm01612986_g1 and TaqMan Fast Advanced Master Mix (Life Technologies), in duplicate, via a Quant Studio 3 qPCR instrument (Thermo Fisher Scientific). The threshold cycle values (Ct) were normalized to the RPL13A housekeeping gene (ΔCt). Relative expressions were calculated via the 2^−ΔCt^ method.

### 2.9. Analysis of cathelicidin in the skin

The level of cathelicidin (CAMP) was measured in the skin homogenates using ELISA kit (Cusabio, CSB-E15061M-96T) according to the manufacturer’s instructions.

### 2.10. Statistical analysis

Descriptive statistics were summarized via standard parametric measures, including the mean and standard deviation (SD). To evaluate changes over time, generalized linear mixed-effects models (GLMMs) with a gamma distribution and a log link function were employed. Both the intercept and slope were treated as random effects for each mouse to account for individual variability. For comparisons between two groups, unpaired t tests were used. One-way analysis of variance (ANOVA) was conducted to compare multiple groups when a single independent variable was involved, followed by post hoc tests where appropriate. Two-way ANOVA was used to assess the effects of two independent variables and their interaction. Post hoc multiple comparisons were performed where appropriate. All the statistical analyses were conducted via R statistical software and GraphPad Prism 8.0.1 (GraphPad Software, Boston, Massachusetts, USA; www.graphpad.com), depending on the specific analysis performed. Graphs were generated in GraphPad Prism 8.0.1. P values *<0.05, **<0.001 and ***<0.0001 were considered significant.

## 3. Results

### 3.1. Determination of MED

The first step in introducing the model was to determine the MED dose for SKH-1 mice. Four different doses (25, 50, 75, and 100 mJ/cm²) were tested and compared with the nonirradiated skin of these mice at each time point (Fig. 1A). None of these doses induced macroscopic effects observable by the naked eye after 24 hours (Fig. 1B). After 48 hours, doses of 25 and 50 mJ/cm² produced noticeable reddening, whereas doses of 75 and 100 mJ/cm² led to the appearance of fine blisters. By 72 hours, fine scabs resembling crusts were observed at the 50 mJ/cm² dose, and areas treated with 75 and 100 mJ/cm² began to peel (Fig. 1B). Visual assessments were complemented by Mexameter-based skin color measurements, which revealed the presence of detectable erythema under all applied conditions at all time points, except in the case of irradiation at 75 and 100 mJ/cm² after 24 h (Fig. 1C). The melanin index of the skin, despite SKH-1 lacking functional melanocytes, did not change noticeably by most of the selected doses (Fig. 1D). In addition, the disruption of the skin’s natural barrier (TEWL) and the hydration of the SC were altered under all tested conditions (Fig. 1E and F). Therefore, 25 mJ/cm² was selected as the MED for subsequent chronic irradiation, as it induced mild erythema by 48 h without overt skin damage (Fig. 1C). The higher doses were deemed excessive and unsuitable for repeated, long-term photoaging.

**Fig. 1:**
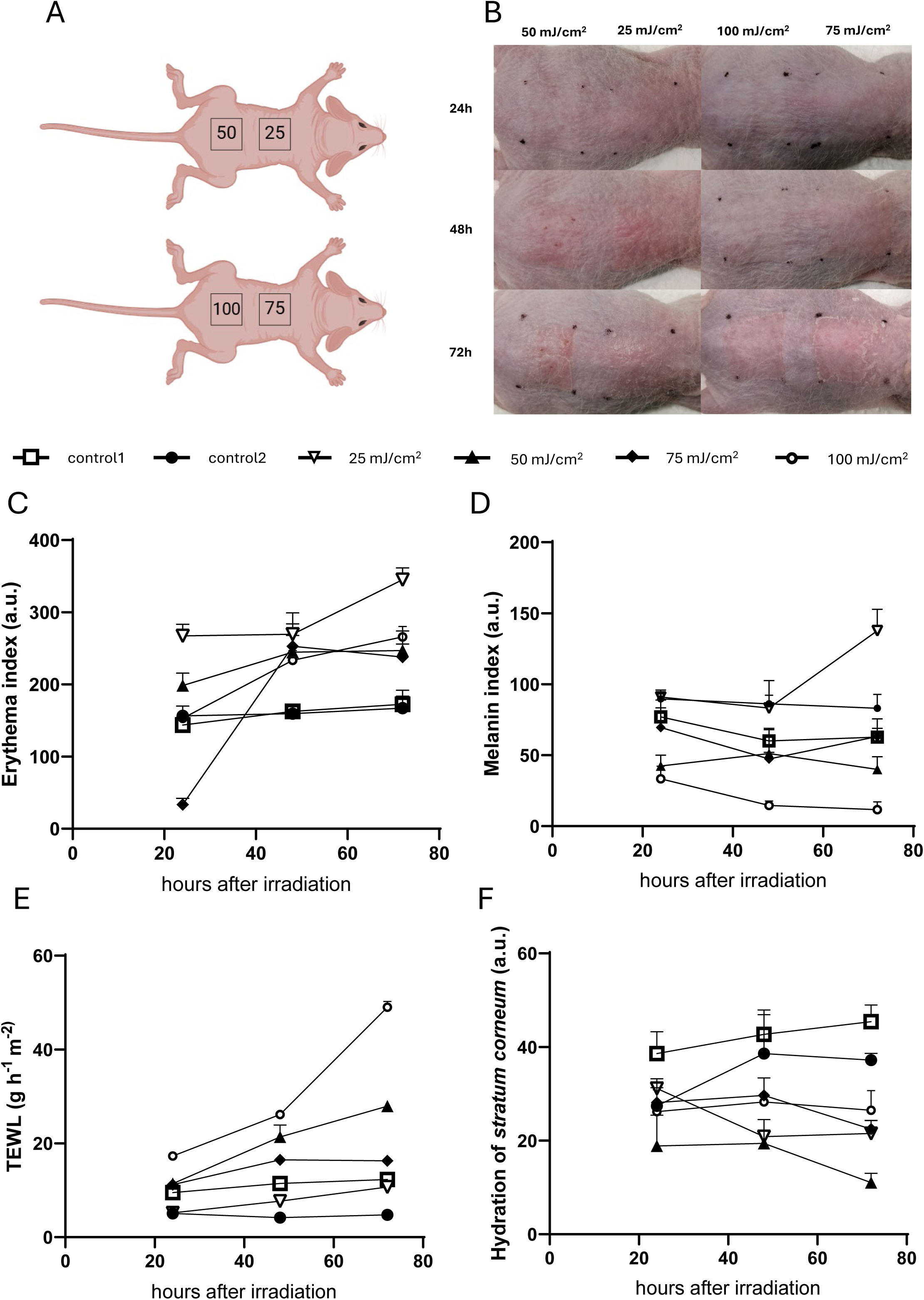
Identification of the optimal UV dose through determination of the MED value. (A) Mice were irradiated with single UV doses of 25, 50, 75, and 100 mJ/cm² to determine MED for SKH-1 mice. (B) Macroscopic evaluation was performed at 24, 48, and 72 hours after UV exposure. (C-F) Quantitative measurements were conducted for the (C) erythema index, (D) melanin index, (E) transepidermal water loss (TEWL), and (F) hydration of the *stratum corneum* of mouse backs irradiated with different UV doses after 24, 48, and 72 hours using MultiProbe Adapter System MPA 580 (Courage + Khazaka electronic GmbH, Germany). Data represent mean values ± standard deviation (*n = 5-7*).

### 3.2. Biphasic timeline of erythema onset and melanogenesis

The manifestation of erythema indicates increased blood flow in the skin and reflects the inflammatory reaction. Thus, to induce photoaging, the skin of SKH-1 mice was irradiated three times a week for a total duration of 2, 4, 6, or 8 weeks (Fig. 2A). The irradiance started at 1 MED and gradually increased to 4 MED over the first 4 weeks, after which the 4 MED levels were maintained for the remainder of the experiment. Although visual skin tone varied slightly among individual SKH-1 mice, objective baseline measurements confirmed no statistical difference between groups prior to irradiation (Fig. 2C and D). A significant effect of irradiation on the erythema index was demonstrated, with a significant interaction between time and treatment (Fig. 2B and C). Interestingly, changes in the erythema index of irradiated skin can be divided into an early response (weeks 0–3) and a late response (weeks 3–8).

**Fig. 2:**
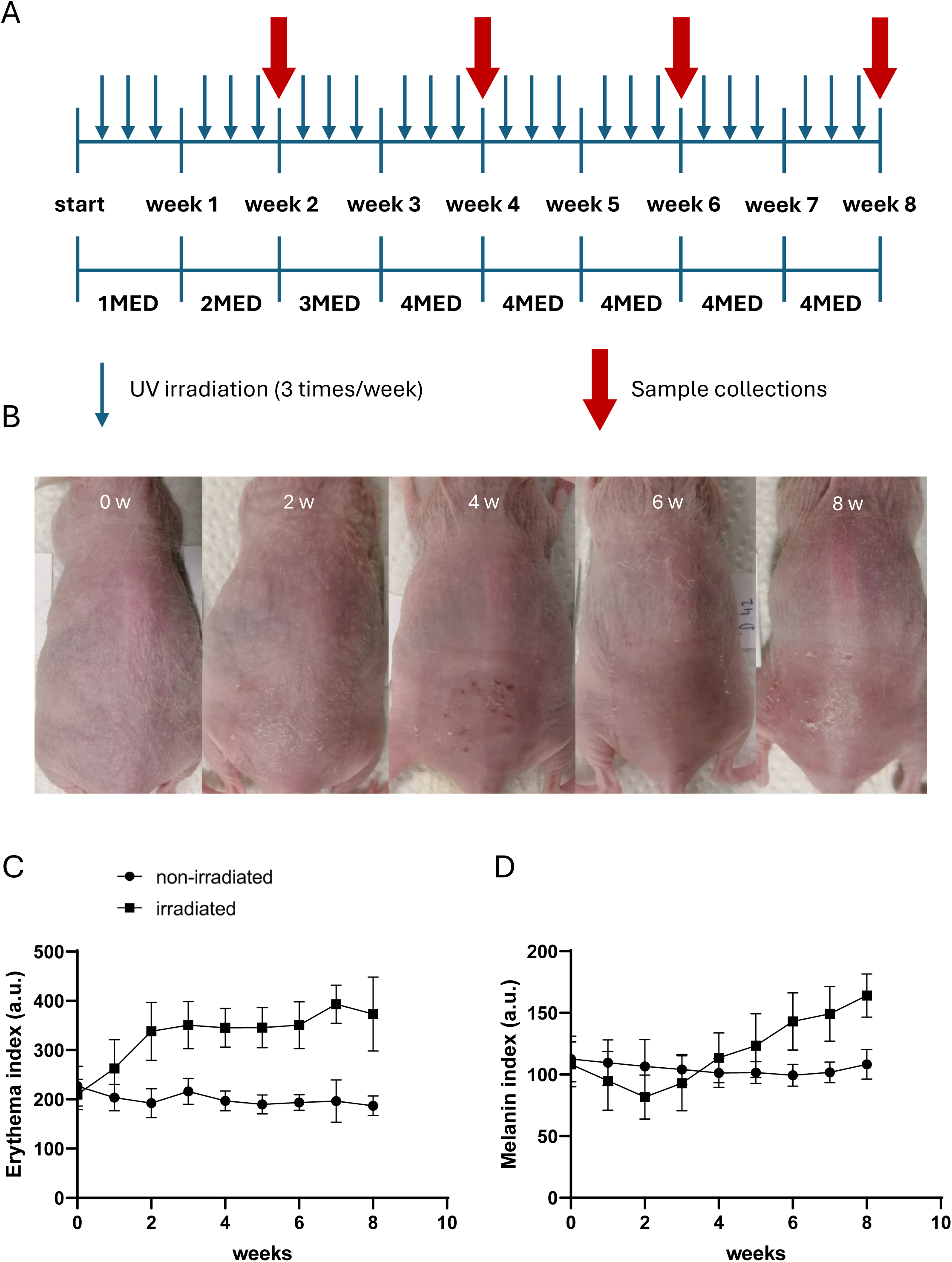
Experimental timeline and skin responses to chronic UV exposure. (A) SKH-1 mice received UV irradiation three times per week with gradually increasing doses from 1 to 4 MEDs over an 8-week period. (B) Representative photographs of these mice were taken to monitor visible skin changes due to UV radiation prior to irradiation and after 2, 4, 6, and 8 weeks of irradiation. (C, D) Erythema and melanin indices were recorded once a week throughout the experiment using the MultiProbe Adapter System MPA 580 (Courage + Khazaka electronic GmbH, Germany). Data represent mean values ± standard deviation (*n = 5-7*).

During the early period, the values in the irradiated group significantly increased, with an average increase of 22% per week, whereas they slightly decreased in the nonirradiated group (approx. 4% per week, exp(β) = 0.96, p = 0.037). From the third week onward, the values in the irradiated group remained stable and were significantly higher (45%) than those in the nonirradiated group (p < 0.001). In the nonirradiated group, the erythema values continued to decrease by an average of 3% per week (p < 0.001). The difference in the trends between the irradiated and nonirradiated groups was statistically significant (p < 0.001). The presence of erythema was also visible macroscopically (Fig. 2B). The melanin index of the skin unexpectedly exhibited two opposite phases when it initially decreased over the first 2 weeks and then increased in the late phase by week 8 (p < 0.05). The values for control nonirradiated mice stagnated (exp(β) = 0.97, p = 0.315), and the difference between the tested groups was significant (p = 0.002) (Fig. 2D).

### 3.3. Skin cell damage and the inflammatory response

Photoaging is associated with chronic inflammation in the dermis as a consequence of UV-mediated damage to cells in the epidermis. Compared with that of nonirradiated control mice, the level of γH2AX, which reflects the degree of DNA damage induced by UV radiation and the formation of DNA double-strand breaks, increased after 2 weeks of irradiation (p = 0.027) and remained significantly greater until the last sampling time at 6 weeks of irradiation (p = 0.002) (Fig. 3A and 4). In control mice, the level of γH2AX was low, indicating only minimal DNA damage in the absence of UV radiation. The epidermal damage induced by chronic UV irradiation was accompanied by a significant increase in the expression of IL-1β (p =0.036) (Fig. 3C). This effect was paralleled by robust neutrophil infiltration into the irradiated dermis (Fig. 5), which was significant compared with that of nonirradiated controls at weeks 6 and 8 (p values of 0.025 and < 0.0001, respectively) (Fig. 3B). Moreover, increased inflammatory activity was confirmed by significantly increased levels of the proinflammatory eicosanoids 6-keto-PGF□α, TXB□, PGF□α, PGE□, PGD□, 12-HETE, and 15-HETE (p values of 0.014, <0.001, 0.001, 0.002, 0.003, <0.0001 and <0.001, respectively) in irradiated skin compared with control skin at 8 weeks (Fig. 3D - J). The level of 5-HETE was not detected in any sample (data not shown).

**Fig. 3:**
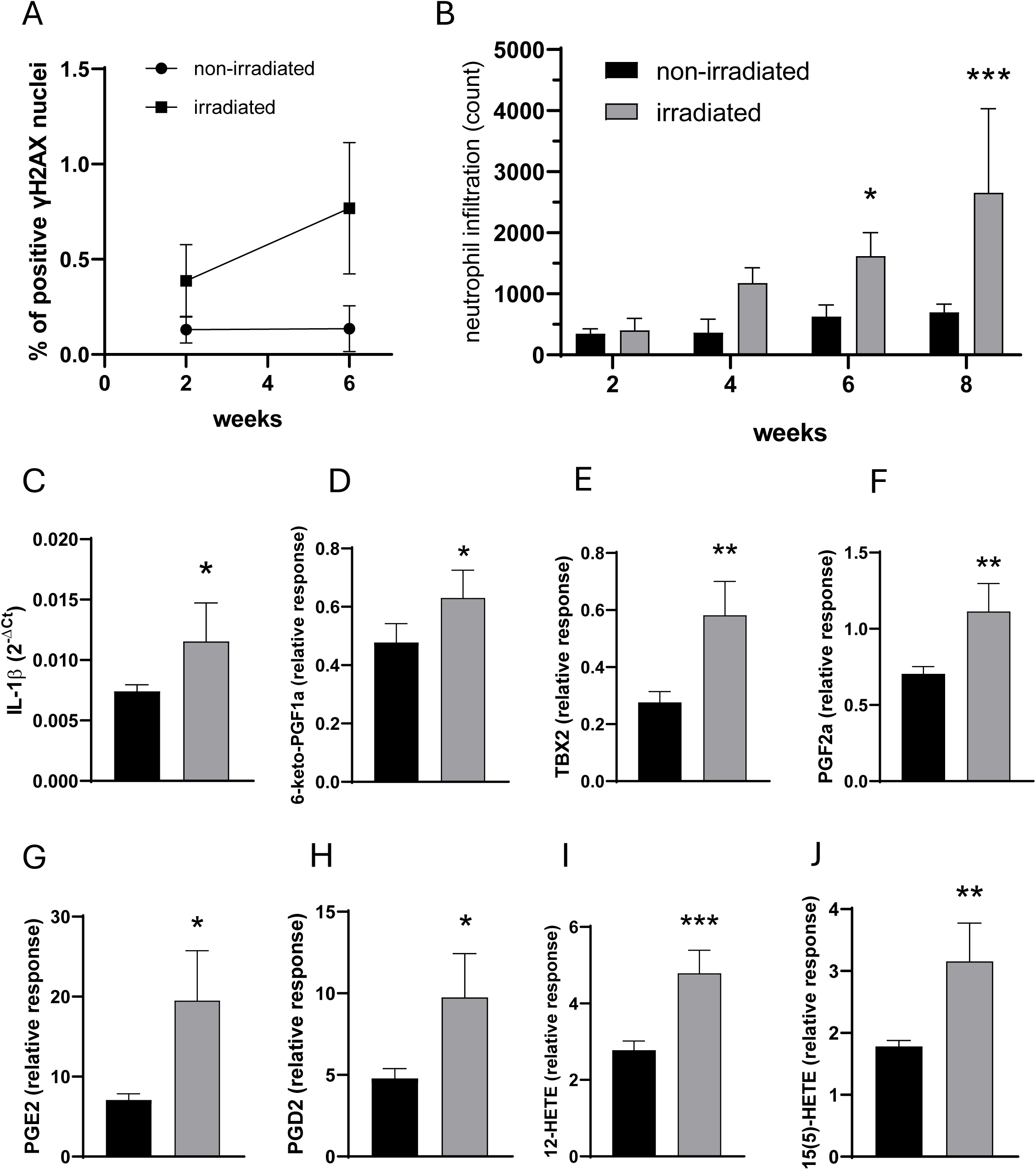
Skin cellular damage and inflammatory response induced by chronic UV exposure. (A) γH2AX in the skin of SKH-1 mice was detected to assess DNA damage after two and six weeks of irradiation. (B) Neutrophil infiltration in the dermis of the mice after 2, 4, 6 and 8 weeks was quantified by immunofluorescence staining. (C) Gene expression of IL-1β in the dermis of these mice was analyzed after 8 weeks of irradiation. (D–J) The levels of selected eicosanoids in the skin of SKH-1 mice after 8 weeks of irradiation were evaluated via lipidomic analysis. Data represent mean values ± standard deviation (*n = 5-7*) (*\* p < 0.05, ** p < 0.001, *** p < 0.0001* compared to nonirradiated control).

**Fig. 4:**
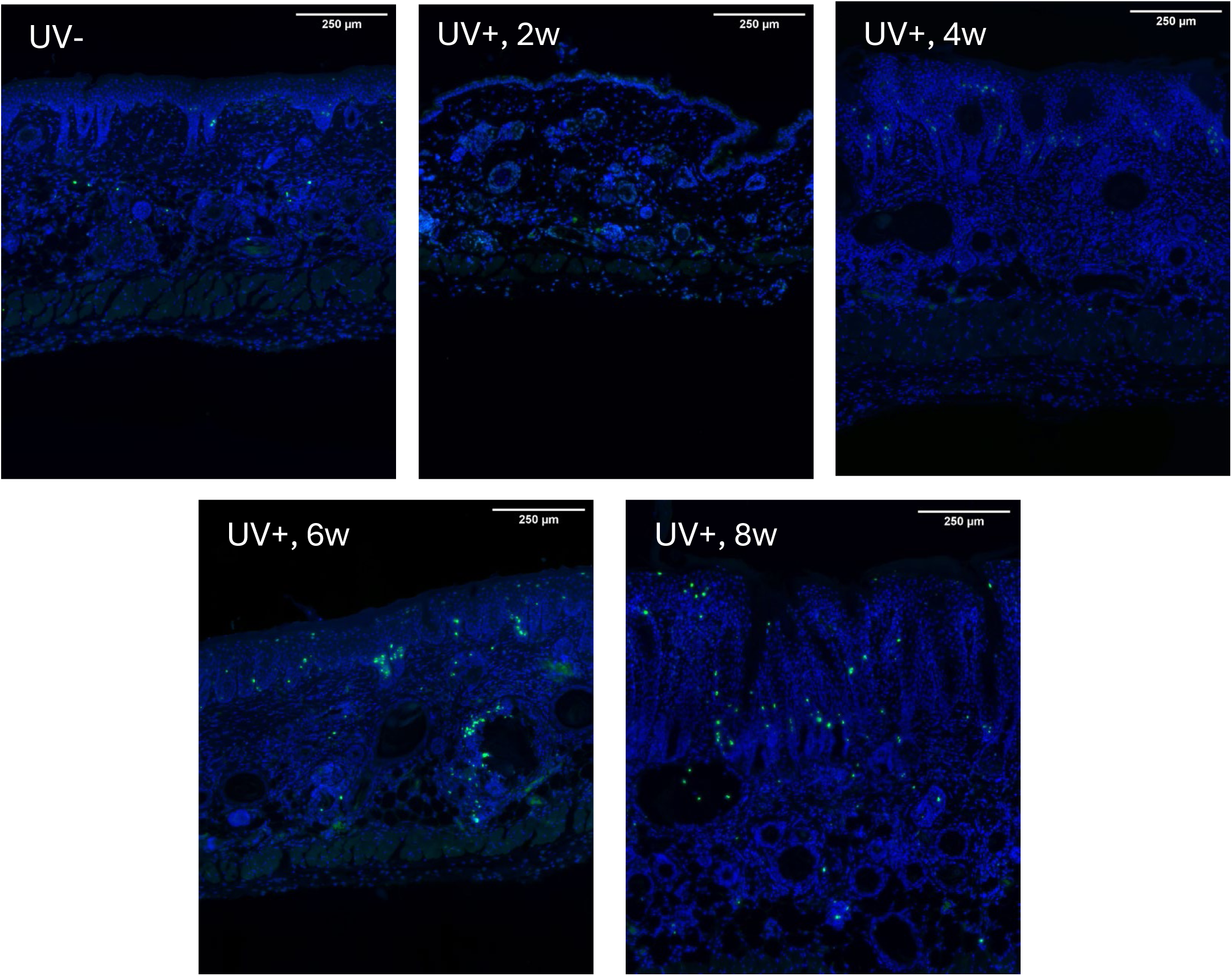
Skin DNA damage induced by UV radiation. γH2AX immunofluorescence staining in the skin sections from nonirradiated and irradiated mice at weeks 2, 4, 6, and 8 was performed to assess DNA damage, as evidenced by a green signal. The cell nuclei are blue, the scale bars at the top of the figures represent 250 μm.

**Fig. 5:**
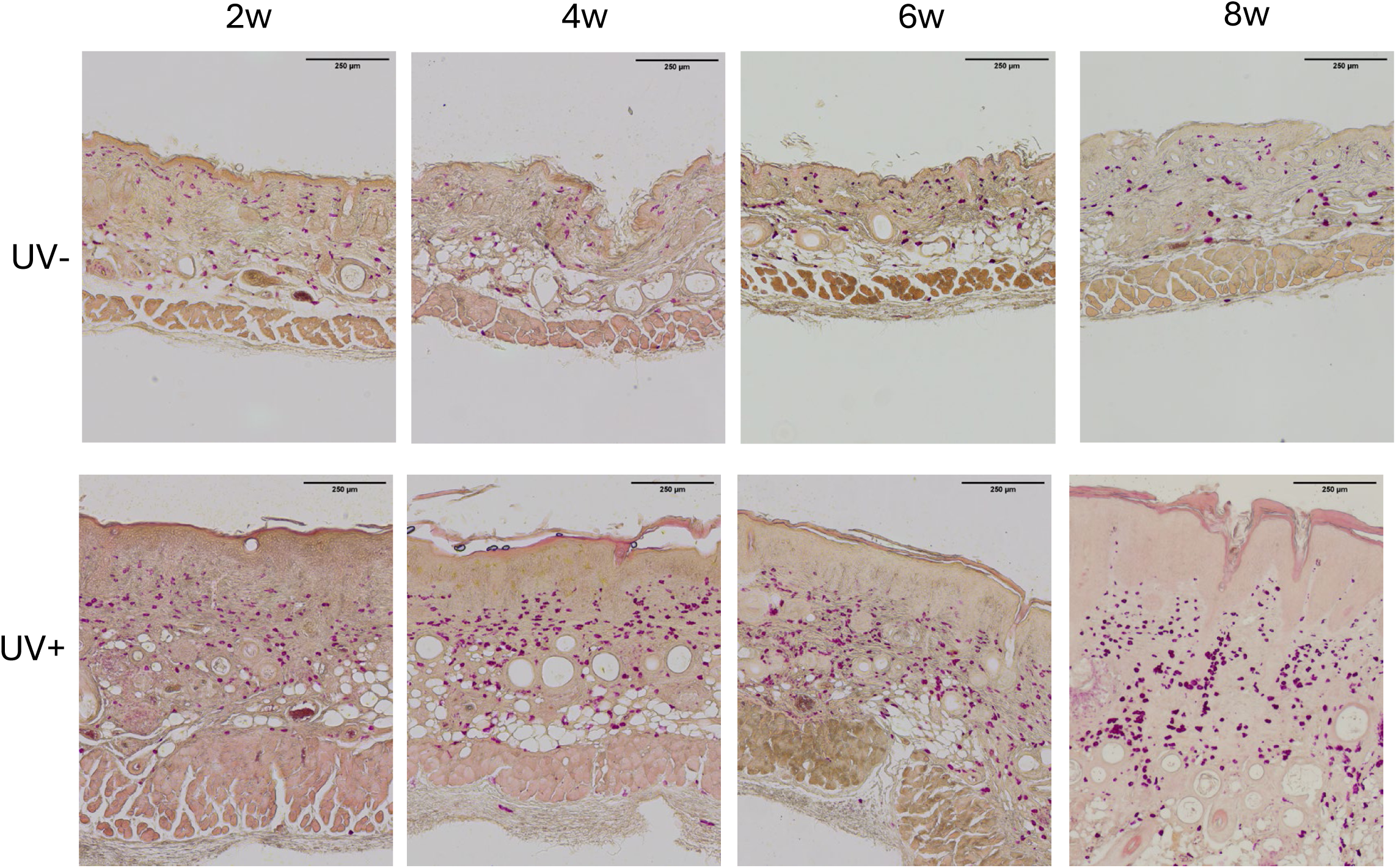
Neutrophile infiltration in nonirradiated (UV-) and irradiated (UV+) skin. Neutrophils were stained with Naphthol AS-D chloroacetate esterase in the mouse skin sections at 2, 4, 6, and 8 weeks. The neutrophils are purple, the scale bar at the top of the figures represents 250 μm.

### 3.4. Skin damage and barrier function impairment caused by UV radiation

Photoaging is related to disruption of the natural barrier of the skin, as indicated by a significant increase in the TEWL, with a significant interaction between time and treatment (<0.001) (Fig. 6A). Although TEWL slightly differed among individual SKH-1 mice, objective baseline measurements confirmed no statistical difference between groups prior to irradiation (Fig. 6A). Interestingly, UV-mediated continuous changes in TEWL can be divided into two phases: early response to irradiation (weeks 0–2) and late response (weeks 2–8). The irradiated mice exhibited a significant increase, with the TEWL increasing by an average of 2.5-fold each week during the first two weeks (exp(β) = 2.45, p < 0.001). From the second week onward, the effect of irradiation reached maximum TEWL levels and remained oscillating around these values, being approximately sixfold greater than that in the nonirradiated group (p < 0.001). In the nonirradiated group, the TEWL values decreased by an average of 8% per week (p = 0.008) after week 2 as the mouse skin physiologically developed with age. Similarly, substantial changes in the hydration of *SC* can also be interpreted in two phases: early response to irradiation (weeks 0–3) and late response (weeks 3–8), with a significant interaction between time and treatment (<0.001) (Fig. 6B). While the values in the nonirradiated group remained stable (exp(β) = 0.97, p = 0.592), those in the irradiated group significantly decreased, with an average decrease of 36% per week over the first three weeks. The impact of photoaging on the hydration of SC then stabilized. However, the values in the irradiated group were significantly lower (approximately 78%) than those in the nonirradiated group (p < 0.001). In both the nonirradiated and irradiated groups, the values increased by an average of 4% per week (p < 0.001). Changes in the TEWL and hydration of the *SC* were also accompanied by a significant increase in epidermal thickness in irradiated mice compared with those in control mice, as shown by histological analysis (Fig. 7). The weekly increase was 36% in the irradiated group compared with 9% in the control group (p = 0.002) (Fig. 8).

**Fig. 6:**
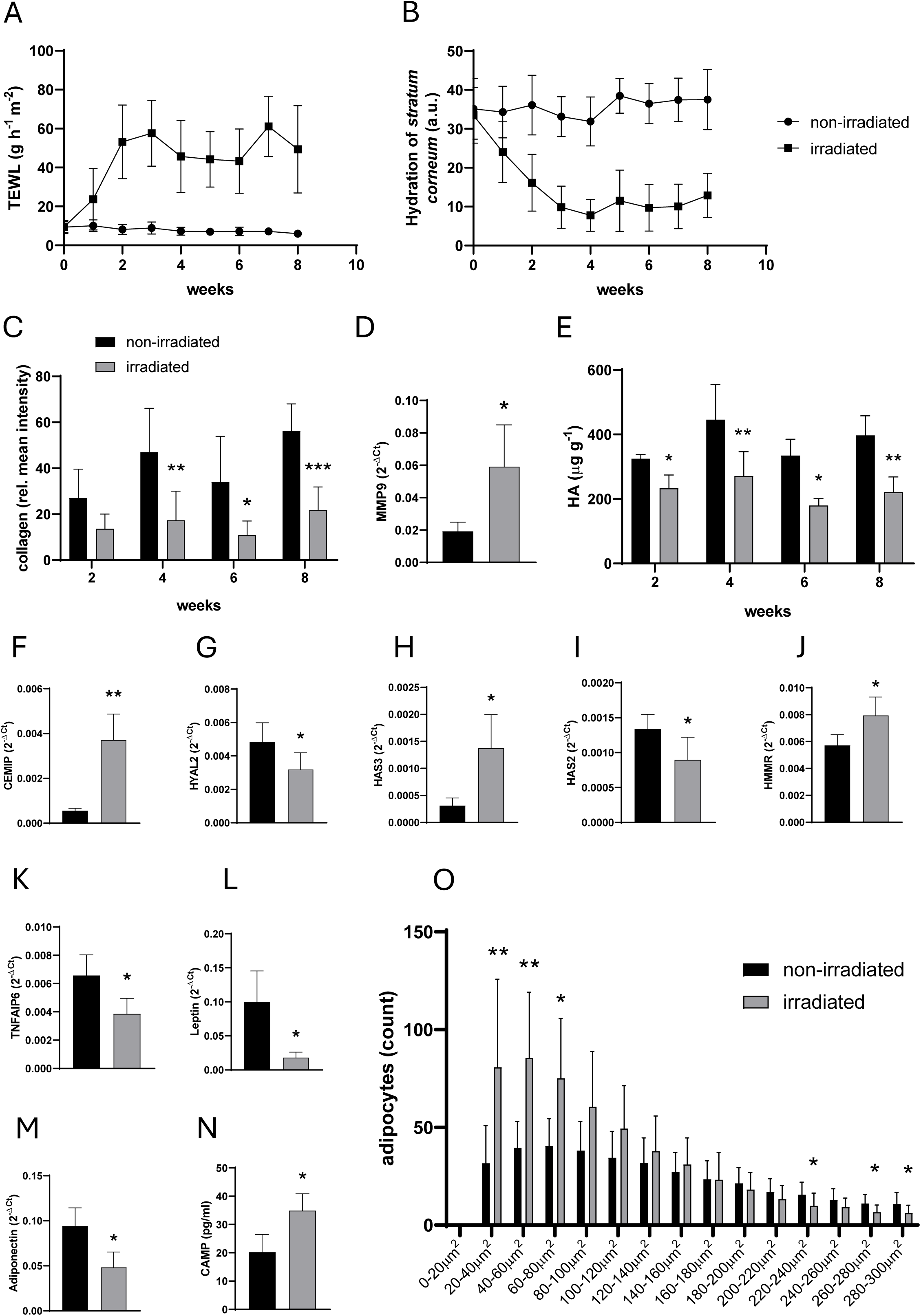
UV-induced skin damage and remodeling of the ECM and adipocytes. (A) TEWL and (B) hydration of the *SC* of SKH-1 mice were recorded once a week during the entire experiment via the MultiProbe Adapter System MPA 580 (Courage + Khazaka electronic GmbH, Germany). (C) Collagen content in the irradiated and nonirradiated mice was quantified at weeks 2, 4, 6, and 8 via histological analysis. (D) MMP9 expression was analyzed in the skin of nonirradiated and irradiated mice after 8 weeks of irradiation. (E) Hyaluronic acid (HA) levels were determined at weeks 2, 4, 6, and 8 via HPLC-MS. (F-K) Gene expression analysis of selected HA-related markers was performed, (L-N) the gene expression of leptin, adiponectin, and cathelicidin was evaluated, (O) adipocyte size distribution was assessed via histological analysis in the skin of nonirradiated and irradiated mice after 8 weeks of irradiation. Data represent mean values ± standard deviation (*n = 5-7*) (*\* p < 0.05, ** p < 0.001, *** p < 0.0001* compared to nonirradiated control).

**Fig. 7:**
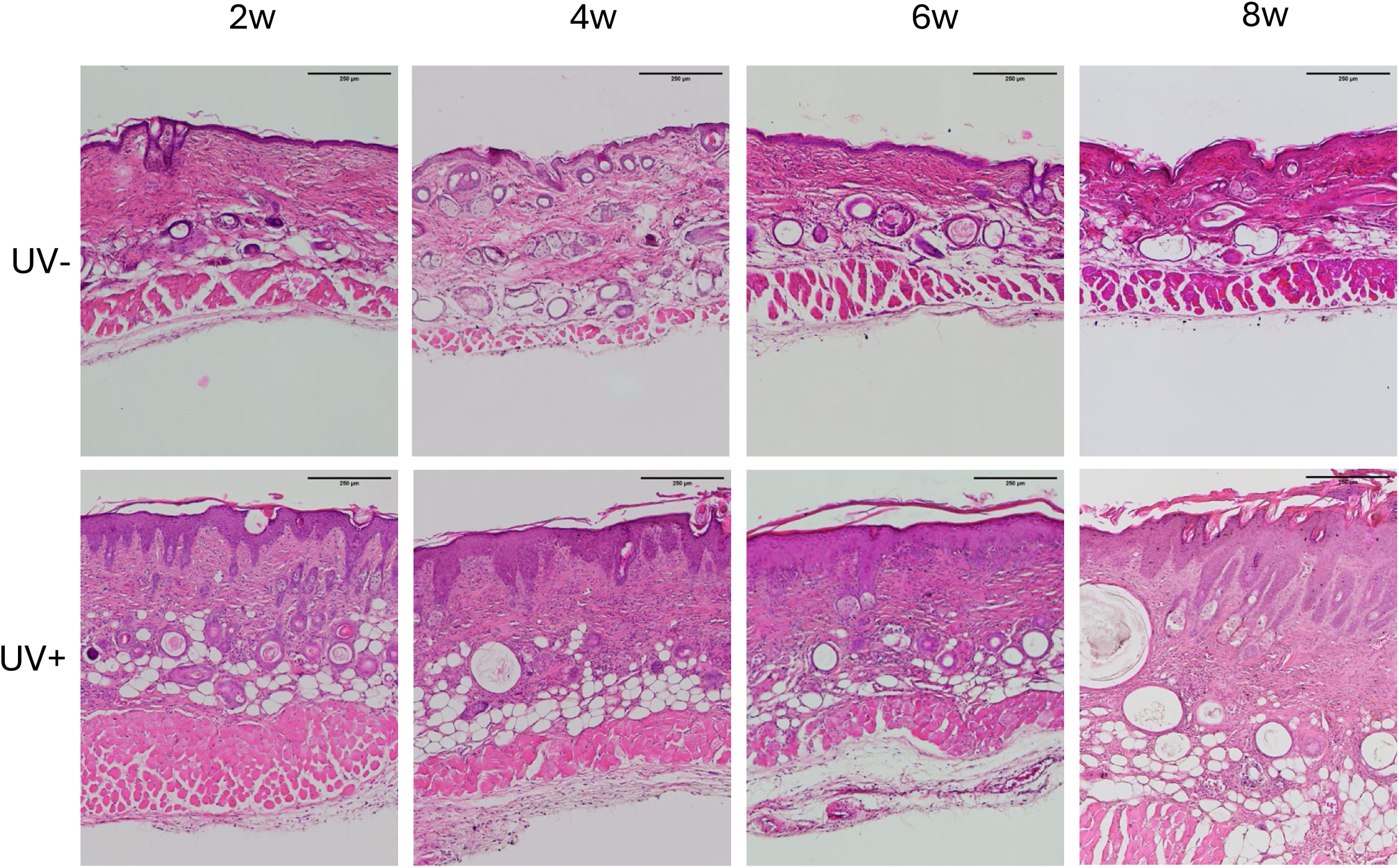
Epidermal thickness in nonirradiated (UV-) and irradiated (UV+) skin. Skin sections from nonirradiated and irradiated mice at weeks 2, 4, 6, and 8 were stained with hematoxylin and eosin to visualize and evaluate epidermal thickness. The scale bar at the top of the figures represents 250 μm.

**Fig. 8:**
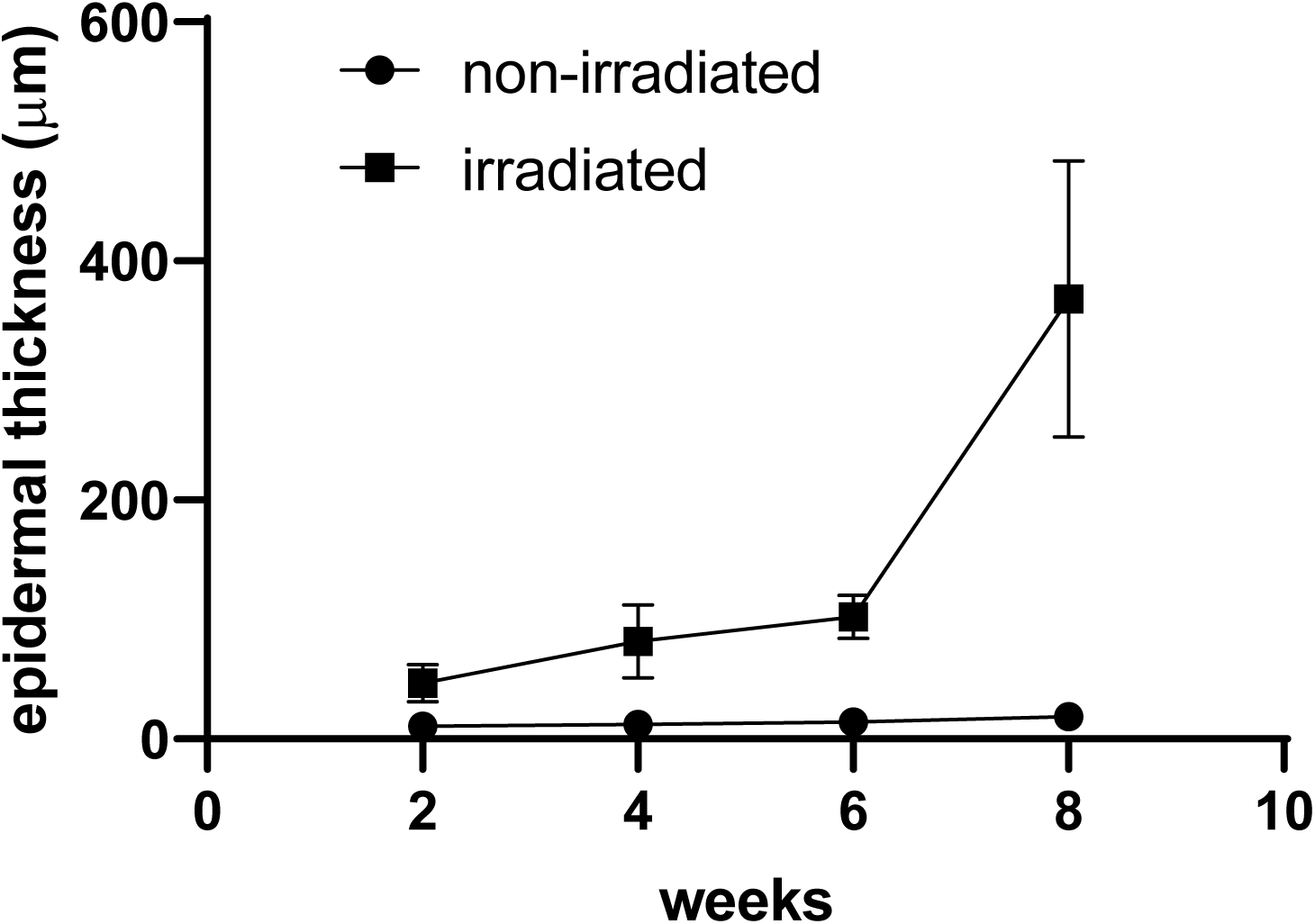
Epidermal thickness increases in irradiated skin. Epidermal thickness of the hematoxylin and eosin-stained skin histological sections from nonirradiated and irradiated mice at weeks 2, 4, 6, and 8 was measured via ImageJ software.

### 3.5. Changes in the transcriptional activity of ECM components and adipocytes

Photoaging was accompanied by substantial changes in the ECM, with the analysis focused on two major components, collagen and HA. Collagen levels in the skin were significantly lower in irradiated mice than in nonirradiated control mice after 4, 6 and 8 weeks of irradiation (p values were <0.001, 0.013 and <0.0001) (Fig. 6C), as shown by histological staining with Masson’s trichrome (Fig. 9). This decrease was accompanied by a significant increase in the gene expression of the major collagen-degrading enzyme MMP9 (p=0.017) (Fig. 6D). Furthermore, the HA content in the skin was significantly lower in irradiated mice than in control mice at every time point (p values at weeks 2, 4, 6 and 8 were 0.01, 0.001, 0.008 and <0.001, respectively) (Fig. 6E). This effect was accompanied by significantly increased expression of the HA-degrading enzyme CEMIP (also known as KIAA1199) (p<0.001) (Fig. 6F) and, in contrast, significant downregulation of HYAL2 (p=0.041) (Fig. 6G). The expression of other detectable HA-degrading enzymes, HYAL1 and CEMIP2 (also known as TMEM2), was not significantly affected (data not shown). The expression of HAS3 enzymes was significantly increased (p=0.011) (Fig. 6H), whereas the expression of HAS2 was significantly decreased (p=0.042) (Fig. 6I). Among HA receptors, CD44 levels remained unchanged (data not shown), whereas RHAMM (protein coded by HMMR gene) was significantly upregulated (p=0.019) in irradiated skin (Fig. 6J). Importantly, TSG-6 expression (protein coded by TNFAIP6 gene) was significantly lower in irradiated skin than in control skin at 8 weeks (p=0.01) (Fig. 6K).

**Fig. 9:**
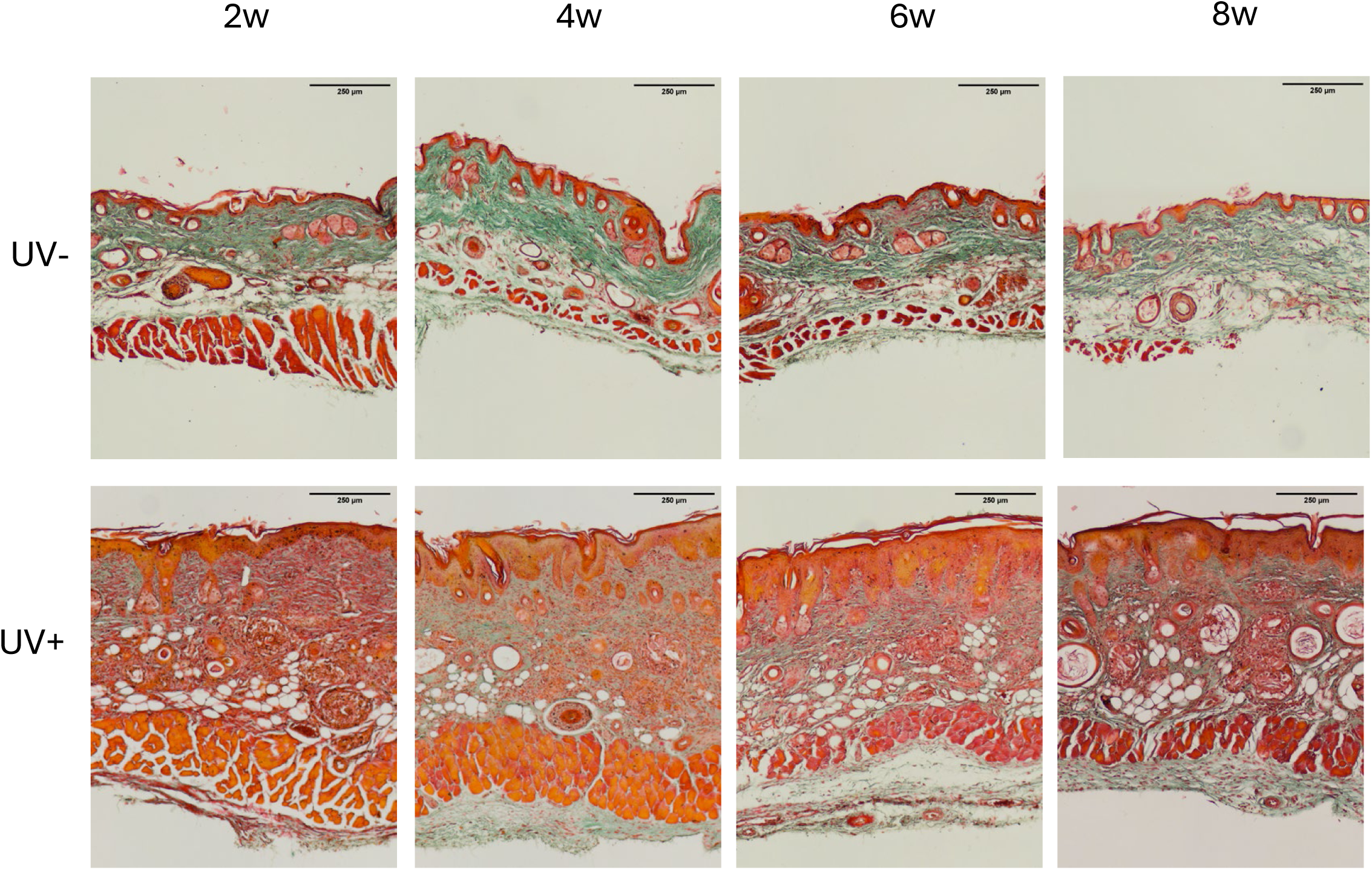
Abundance and collagen density in nonirradiated (UV-) and irradiated (UV+) skin. Collagen fibrils were visualized via Masson-trichrome staining and appear green in the histological skin sections from nonirradiated and irradiated mice at weeks 2, 4, 6, and 8. The scale bar at the top of the figures represents 250 μm.

Finally, photoaging-driven changes in subcutaneous fat indicate an active role of adipocytes in inflammation and in UV-irradiated skin. Accordingly, the gene expression of two typical peptides produced by adipocytes, leptin and adiponectin (Fig. 6L and M), was significantly decreased in the irradiated skin (p values of 0.002 and 0.005, respectively). Interestingly, there was an increase in the concentration of the antimicrobial peptide cathelicidin in UV-exposed skin (p=0.002) (Fig. 6N). Histological analysis of mouse skin samples revealed significant differences in the number and size distribution of adipocytes between UV-irradiated and nonirradiated mice at all the observed time points (2, 4, 6, and 8 weeks) (p < 0.001) (Fig. 6O). UV-irradiated mice presented significantly greater numbers of smaller adipocytes in the 20–40 μm² (p < 0.001), 40–60 μm² (p = 0.001), and 60–80 μm² (p = 0.008) size ranges. Conversely, nonirradiated mice had significantly greater numbers of larger adipocytes in the 220–240 μm² (p = 0.022), 260–280 μm² (p = 0.026), and 280–300 μm² (p = 0.016) categories. These findings suggest that UV exposure is associated with a shift toward a population of smaller adipocytes.

## 4. Discussion

In this study, repeated UV irradiation of SKH-1 mice gradually increased to 4 MED, resulting in a characteristic two-phase erythema response, DNA damage (γH2AX), and robust neutrophil infiltration accompanied by elevated IL-1β expression, which is a hallmark of the inflammatory process that drives photoaging. The barrier function of the skin was significantly disrupted, as demonstrated by increased TEWL, reduced SC hydration, and epidermal thickening. The collagen content tended to decrease, and HA was significantly depleted, coupled with imbalances in enzymes regulating these matrix components. Furthermore, the loss of subcutaneous fat was indicated by altered expression of adipokines, leptin and adiponectin, and histological evidence of smaller adipocytes. This study not only documents multiple UV-induced alterations but also determines how these parameters evolve over time and which of them are most informative at specific stages of photoaging. The broad panel of endpoints analyzed in this work, therefore, defines a temporal hierarchy of indicators that characterize early, transitional, and late phases of chronic UV-induced skin damage. This time-resolved perspective is essential for selecting appropriate readouts in future studies using this model.In this work, 25 mJ/cm² (1 MED) was selected for chronic exposure, the lowest dose at which erythema appeared on the skin of SKH-1 mice within 24–48 hours. There is a significant variation in MED values reported in previous studies using mouse models to investigate the effects of UV radiation on the skin, ranging from approximately 40 to 90 mJ/cm².^25–28^ These variations can be attributed to differences in experimental protocols, such as the use of different UV radiation sources or filters, as well as variations in mouse strains, sex, or age. However, these findings are consistent with those reported in humans, in which skin pigmentation, thickness, body region, and previous UV exposure influence individual skin sensitivity.^29^ Authors Gyongyosi et al.^25^ compared acute skin reactions to UVB radiation among 3 mouse strains, SKH-1, C57BL/6 N, and BALB/c, with the mice being irradiated and euthanized after 24 and 48 hours. These findings suggest that the reaction of the skin of the C57BL/6 N mouse strain was the most similar to the reaction of human skin because of the presence of melanin pigmentation. However, C57BL/6 N mice need to be shaved 24 hours before irradiation, which can cause greater irritation to the skin than UV irradiation and generate artifact data. Unlike most studies focusing on a single endpoint, we monitored sequential changes, providing a more complete understanding of UV-induced skin processes over time.

Interestingly, biphasic erythema development was observed in our study. Erythema revealed sustained elevation after an initial rapid increase. This observation matches the suggested progression of photoaging through early and late phases, early transient vs. chronic persistent inflammation, each of which is characterized by distinct biological events. As discussed in prior studies, in the initial phase, UV-driven oxidative stress and acute inflammation cause vasodilation of cutaneous blood vessels and rapidly increase erythema.^30,31^ The persistent inflammation subsequently mediates the transition to a late phase dominated by extensive remodeling of the dermal ECM (discussed below). Although SKH-1 mice are nonpigmented and lack functional melanocytes capable of melanogenesis,^32^ the observed changes in the melanin index align with UV-induced processes affecting skin properties. The initial decrease in the melanin index during the first two weeks may be attributed to mild desquamation observed during this period, which likely alters the skin’s optical properties. The subsequent late-phase increase in the melanin index matches the elevated erythema levels. This finding suggests that other UV-induced skin adaptations, such as epidermal thickening and increased blood flow, contribute to changes in optical properties. Baquié and Kasraee (2014) compared two skin colorimeters, Dermacatch® and Mexameter®, and reported that the melanin and erythema values of the Mexameter were falsely affected by increases in erythema or variations in pigmentation, respectively.^33^ One key factor influencing these readings is similar absorption spectra of hemoglobin and melanin ^34^ Thus, the observed fluctuations in the melanin index in SKH-1 mice are likely a result of UV-induced modifications in epidermal thickness, hemoglobin concentration, and overall optical skin properties rather than genuine melanogenesis. When interpreting pigmentation measurements in nonpigmented models via reflectance-based devices, these factors should be carefully considered. The increase in the erythema index in irradiated mice may indicate that inflammatory processes occur in the skin, particularly in the epidermal layer. The inflammatory process is induced by UV-mediated damage to cells. Although γH2AX is not yet a routine marker in photoaging studies, it is a sensitive indicator of UV-induced DNA double-strand breaks.^35^ The increased number of γH2AX foci in irradiated skin confirmed the accumulation of DNA damage in our model. There is a dose-dependent relationship between UV exposure and DNA damage, as reflected by the accumulation of γH2AX foci. The highest level of γH2AX was observed after 6 weeks of UV irradiation, which indicates that the peak level of DNA damage occurred in response to UV exposure.

Continued DNA damage highlights the negative effects of UV radiation on cellular integrity, confirming its role in inducing genetic stress and triggering limited repair responses, resulting in a potent inflammatory stimulus in photodamaged skin. Taken together, these observations indicate that erythema, TEWL, SC hydration, and γH2AX, all of which were used to assess UV-induced changes in our study, represent sensitive indicators of the early inflammatory and barrier-disruption phase of photoaging (weeks 0–3), when functional skin impairment precedes substantial structural remodeling. These parameters are therefore particularly suitable for studies focused on acute or early UV responses.

The intense inflammatory process is characterized by significantly increased expression of IL-1β, which is primarily induced by the intracellular signaling machinery involved in inflammasome formation. The production of IL-1β-mediated ongoing inflammatory processes is consistent with the observed neutrophil influx. The increase in the leukocyte infiltration corresponds with observations in the study by Wang et al.,^36^ where nude mice were irradiated by UVA radiation. The inflammation process is connected with the infiltration of phagocytes, which release a wide range of degrading enzymes, and is one of the key factors in the massive effects of photoaging on the ECM. The undergoing inflammatory process is further documented by a significant increase in proinflammatory eicosanoids in chronically UV-irradiated skin and reflects the activation of arachidonic and linoleic acid metabolism pathways, particularly cyclooxygenase and lipoxygenase enzymes. Prostaglandins such as PGE□ and PGD□, products of cyclooxygenase pathways, are implicated in different inflammatory conditions, promoting vasodilation, erythema, and immune cell infiltration.^37^ Similarly, TXB□ contributes to local inflammatory responses through vasoconstriction and modulation of inflammatory cell recruitment.^37^ Moreover, increased 12-HETE and 15-HETE levels suggest activation of the lipoxygenase pathways, known for their roles in neutrophil chemotaxis and exacerbating inflammatory responses.^37^

The persistent inflammatory milieu and sustained activation of signaling pathways mediate the transition to a late phase dominated by extensive remodeling of the dermal ECM. During this phase, proteolytic enzymes, such as MMPs, degrade collagen, while altered HA metabolism contributes to the gradual loss of structural integrity and a more chronically inflamed microenvironment. Collagen and HA were chosen for evaluation in our experiment because they are key components of the skin ECM. UV-exposed skin samples consistently presented lower HA concentrations than did their respective nonexposed controls at each time point, reflecting the ability of UV to induce a processes leading to HA degradation. Despite the absence of a significant difference in collagen content between irradiated and control mice, the values were consistently lower in irradiated mice than in control mice across all time points. The observed variability in HA levels among controls and irradiated mice may indicate natural fluctuations in HA metabolism. This was accompanied by significant changes in HA-degrading enzymes, with CEMIP significantly increased and HYAL2 significantly downregulated. The expression of other detectable HA-degrading enzymes, HYAL1 and CEMIP2, was not significantly affected. Similarly, the opposite gene expression pattern was observed for HA-synthesizing enzymes when HAS3 was significantly increased, while the expression of HAS2 significantly decreased. Previously, in mice chronically irradiated with UVB for 182 days, the gene expression levels of HAS1, HAS2, and HAS3 were significantly decreased, whereas those of HYAL1 and HYAL2 were not significantly altered.^38^ This was accompanied by a significant loss of HA from the papillary dermis. In contrast, no significant HA loss after 20 weeks of UVB irradiation in mice was reported by Rock et al.^39^, which could be due to the timing of HA turnover analysis. In our study, the HA receptor CD44 was not modified by chronic irradiation, in contrast to RHAMM, which was significantly increased. Similarly, no significant modification of CD44 in chronically irradiated mice was reported by Dai et al.^38^ TSG-6 expression was significantly lower in irradiated skin than in control skin at 8 weeks. This decrease in the level of TSG-6, a protein with anti-inflammatory and ECM-protective effects, suggests that chronic UV irradiation may impair the natural protective feedback mechanisms of the skin.

Pathological changes in the ECM, decreased SC hydration, and skin barrier disruption manifest the chronic effects of photoaging achieved by long-term skin exposure to UV radiation. Similar findings were reported in a study by Pyun et al.,^40^ where mice irradiated with UVB radiation for up to 12 or 14 weeks presented a significant decrease in skin hydration and a significant increase in TEWL at the study endpoints. Despite the application of different spectra, the results of our 8-week experiment correspond to their observations, as these parameters are mainly characteristic of the epidermis, which is affected by UVB radiation.

When interpreting the values of epidermal thickness, it is important to consider that the model starts with different baseline values for irradiated and nonirradiated mice because the first evaluation of skin thickness was performed 2 weeks after the initiation of the experiment. The observed differences in values could already reflect the effects of irradiation or could be attributed to variations in the initial skin thickness of the mice, despite being randomly selected. Interestingly, after eight weeks of UV radiation, the epidermal thickness significantly increased compared with that at earlier time points. This sharp increase could indicate that severe skin damage may have occurred at this stage, leading to extreme hyperplasia. An increase in the skin thickness, as demonstrated by H&E staining, was also observed in the previously mentioned study by Pyun et al.^40^ This pattern of epidermal thickening, which reflects a proliferative response, indicates that UV exposure drives a dose-dependent increase in skin cell turnover as a reaction to damage.

Studies of changes in subcutaneous adipose tissue in chronic UV-exposed skin have revealed somewhat controversial results. Our results suggest an explanation for these differing findings. We detected a significantly greater number of smaller, immature adipocytes alongside increased antimicrobial peptide cathelicidin and decreased expression of key adipokines, such as leptin and adiponectin, in UV-irradiated skin. Leptin is produced by mature adipocytes, and cathelicidin is released during adipogenesis and attenuates inflammation.^41,42^ The UV-induced inhibition of adipokine production, as described by Kim et al. (2016),^43^ aligns with our findings. Moreover, the levels of these adipokines are critical for maintaining normal skin homeostasis, immune function, and ECM remodeling. The higher levels of cathelicidin and the greater number of immature adipocytes in UV-irradiated skin in our study could support the “reactive” adipogenesis theory.^42^ Liggins et al. (2019) revealed an increase in cathelicidin during reactive adipogenesis, which is a process of inflammation-associated fat remodeling described recently in the context of inflammatory conditions not related to UV.^7,10^ On the other hand, our results simultaneously revealed a reduced number of mature adipocytes following UV exposure. Interestingly, decreased lipogenesis in subcutaneous fat was observed by Kim et al. through the suppression of key adipogenic transcription factors.^8^ These dysregulated adipokines can perpetuate local inflammation, interfere with collagen biosynthesis, and augment degradation processes.^7,9^

In contrast to the early functional changes, alterations in HA metabolism, collagen organization, CEMIP/HAS expression, TSG-6 downregulation, and the shift in adipocyte size distribution become prominent only after prolonged irradiation. These markers, therefore, frame the late remodeling phase of photoaging, characterized by structural extracellular matrix degradation and reactive adipose tissue changes. These endpoints are consequently more appropriate for evaluating interventions targeting chronic tissue remodeling rather than early inflammatory responses.

Comparative studies suggest that the molecular and histopathological features of UV-induced photoaging in SKH-1 mice closely mirror those observed in human skin, supporting the translational relevance of this animal model. Repeated measurements of parameters typical for studies involving human volunteers were applied in this work to support the other parameters frequently analyzed in studies involving *in vivo* mouse models, where the primary focus is typically to examine histological and molecular changes induced by UV radiation.^25,26,44^ One advantage of such a study is the ability to analyze the full thickness of the skin obtained from laboratory animals, whereas in human studies, samples of only the epidermis are collected. Although variations in skin anatomy and immune responses exist between species, key hallmarks of human photoaging—such as disrupted collagen architecture, chronic low-grade inflammation, and impaired barrier function—are consistently observed in murine models exposed to cumulative doses of UV radiation. Additionally, the progressive changes in ECM remodeling in mice reflect similar time-dependent processes described in human photoexposed epidermis and dermis. Notably, our MED falls within the range of human skin type I MED.^45^ By modulating irradiation protocols to approximate habitual sun exposure patterns, researchers can replicate many of the characteristic phenotypes of photoaged human skin, including wrinkling, pigmentary alterations, and loss of elasticity.

Taken together, based on time dependent changes of monitored parameters, three biologically distinct phases of chronic UV-induced photoaging can be distinguished in this model: (i) an early inflammatory/barrier dysfunction phase (up to third week), (ii) a transitional phase marked by epidermal hyperplasia and leukocyte infiltration (weeks 3–4), and (iii) a late phase with significant extracellular matrix and adipose tissue remodeling phase (weeks 4–8). This distinction clarifies which endpoints are most informative depending on the experimental objective and prevents misinterpretation arising from the use of inappropriate markers at incorrect time points. Although these mouse models provide valuable insights into the effects of UV radiation on skin biology, several limitations must be acknowledged. First, species-specific differences exist between murine and human skin, particularly in terms of immune response and anatomy, including thickness, hair follicle density, and melanocyte activity. Next, the duration of the study, while sufficient to observe early and progressive skin changes, may not fully capture long-term adaptations or chronic effects such as sustained ECM remodeling, persistent inflammation, or carcinogenesis. By addressing these model constraints through complementary approaches, future research can bridge the gap between preclinical findings and human applications.

## 5. Conclusion

Chronic UV irradiation in SKH-1 mice induced a phenotype closely resembling human photoaging, marked by sustained inflammation and progressive tissue alterations. We observed a two-phase erythema response and significant barrier dysfunction, along with epidermal hyperplasia, by week 8. UV exposure disrupted the dermal extracellular matrix. HA levels decreased at all time points, and collagen showed signs of degradation. UV exposure profoundly affected subcutaneous fat, reducing adipocyte size and lowering the expression of adipocyte-derived factors, indicating a significant shift in adipocyte development. This coupling of ECM deterioration with “reactive” adipose tissue changes underscores that photoaging is not confined to the epidermis but involves deeper skin layers. Our well-characterized model highlights key molecular targets of photoaging, from DNA damage and inflammatory mediators to matrix-regulating enzymes and protective factors such as TSG-6, whose expression is decreased. These findings deepen our understanding of UV-driven skin aging and point to potential intervention strategies. Consequently, findings from SKH-1 mouse experiments offer critical insights into both the mechanisms underpinning human photoaging and the development of targeted therapeutic strategies.

## 6. Data Availability Statement

The data that support the findings of this study are available from the corresponding author upon reasonable request.

## 7. Author Contribution / Credit statement

Conceptualization: MB, RŠ, MŠ, LK, KN, ID; Data Curation: MB, RŠ, PO, MŠ, MČ; Formal Analysis: MB, RŠ, PO, MŠ, MČ; Funding Acquisition: VV, LK; Investigation: MB, RŠ, MŠ, KL, MČ; Methodology: MB, RŠ, MŠ, KL, ID; Project Administration: LK, KN, VV; Supervision: LK, KN, VV; Visualization: MB, RŠ, KL

## 8. Acknowledgements

This work was supported by the institutional support of the Institute of Biophysics of the Czech Academy of Sciences (68081707) and by the Operational Programme Johannes Amos Comenius, project “Biology of hyaluronic acid” No. CZ.02.01.01/00/23_020/0008499 of the Ministry of Education, Youth and Sports of the Czech Republic.

## List of abbreviations

ANOVA: Analysis of variance
BSA: Bovine serum albumin
CAMP: Cathelicidin antimicrobial peptide
C/EBPα: CCAAT/enhancer-binding protein alpha
CCL5: C-C motif chemokine ligand 5
CCL20: C-C motif chemokine ligand 20
CEMIP: Cell migration-inducing protein
CXCL5: C-X-C motif chemokine ligand 5
DAPI: 4′,6-diamidino-2-phenylindole
DNA: Deoxyribonucleic acid
ECM: Extracellular matrix
GLMM: Generalized linear mixed-effects model
HA: Hyaluronic acid (also referred to as hyaluronan)
HAS: Hyaluronan synthase
H&E: Hematoxylin and eosin
HETE: hydroxyeicosatetraenoic acid
HYAL: Hyaluronidase
IL: Interleukin
MED: Minimal erythema dose
MMP: Matrix metalloproteinase
PPARγ: Peroxisome proliferator-activated receptor gamma
qPCR: Quantitative polymerase chain reaction
RHAMM: Receptor for hyaluronan-mediated motility
RT: Room temperature
SC: Stratum corneum
SD: Standard deviation
SREBP-1: Sterol regulatory element-binding protein 1
TBS: Tris-buffered saline
TEWL: Transepidermal water loss
TNF-α: Tumor necrosis factor alpha
TSG-6: TNFα-induced protein 6
UV: Ultraviolet

## References

1. Farage MA, Miller KW, Elsner P, Maibach HI. Intrinsic and extrinsic factors in skin ageing: a review. International Journal of Cosmetic Science. 2008;30(2):87–95. 10.1111/j.1468-2494.2007.00415.x

2. Ansary TM, Hossain MR, Kamiya K, Komine M, Ohtsuki M. Inflammatory Molecules Associated with Ultraviolet Radiation-Mediated Skin Aging. Int J Mol Sci. 2021;22(8). doi:10.3390/ijms22083974

3. Battie C, Jitsukawa S, Bernerd F, Del Bino S, Marionnet C, Verschoore M. New insights in photoaging, UVA induced damage and skin types. Exp Dermatol. 2014;23 Suppl 1:7–12. doi:10.1111/exd.12388

4. Quan T, He T, Kang S, Voorhees JJ, Fisher GJ. Solar Ultraviolet Irradiation Reduces Collagen in Photoaged Human Skin by Blocking Transforming Growth Factor-β Type II Receptor/Smad Signaling. The American Journal of Pathology. 2004;165(3):741–751. 10.1016/S0002-9440(10)63337-8

5. Fisher GJ, Wang ZQ, Datta SC, Varani J, Kang S, Voorhees JJ. Pathophysiology of premature skin aging induced by ultraviolet light. N Engl J Med. 1997;337(20):1419–1428. doi:10.1056/nejm199711133372003

6. Kimball AB, Alora-Palli MB, Tamura M, et al. Age-induced and photoinduced changes in gene expression profiles in facial skin of Caucasian females across 6 decades of age. Journal of the American Academy of Dermatology. 2018;78(1):29–39.e7. 10.1016/j.jaad.2017.09.012

7. Kim EJ, Kim YK, Kim S, et al. Adipochemokines induced by ultraviolet irradiation contribute to impaired fat metabolism in subcutaneous fat cells. British Journal of Dermatology. 2018;178(2):492–501. doi:10.1111/bjd.15907

8. Kim EJ, Kim YK, Kim JE, et al. UV Modulation of Subcutaneous Fat Metabolism. Journal of Investigative Dermatology. 2011;131(8):1720–1726. doi:10.1038/jid.2011.106

9. Lee J, Lee J, Jung E, et al. Ultraviolet A Regulates Adipogenic Differentiation of Human Adipose Tissue-derived Mesenchymal Stem Cells via Up-regulation of Kruppel-like Factor 2. Journal of Biological Chemistry. 2010;285(42):32647–32656. doi:10.1074/jbc.M110.135830

10. Liggins MC, Li F, Zhang L juan, Dokoshi T, Gallo RL. Retinoids Enhance the Expression of Cathelicidin Antimicrobial Peptide during Reactive Dermal Adipogenesis. The Journal of Immunology. 2019;203(6):1589–1597. doi:10.4049/jimmunol.1900520

11. Moloney SJ, Edmonds SH, Giddens LD, Learn DB. The hairless mouse model of photoaging: evaluation of the relationship between dermal elastin, collagen, skin thickness and wrinkles. Photochem Photobiol. 1992;56(4):505–511. doi:10.1111/j.1751-1097.1992.tb02194.x

12. Benavides F, Oberyszyn TM, VanBuskirk AM, Reeve VE, Kusewitt DF. The hairless mouse in skin research. Journal of Dermatological Science. 2009;53(1):10–18. 10.1016/j.jdermsci.2008.08.012

13. Cachon-Gonzalez MB, Fenner S, Coffin JM, Moran C, Best S, Stoye JP. Structure and expression of the hairless gene of mice. Proceedings of the National Academy of Sciences. 1994;91(16):7717–7721. doi:doi:10.1073/pnas.91.16.7717

14. Gong M, Zhang P, Li C, Ma X, Yang D. Protective Mechanism of Adipose-Derived Stem Cells in Remodelling of the Skin Stem Cell Niche During Photoaging. Cell Physiol Biochem. 2018;51(5):2456–2471. doi:10.1159/000495902

15. Chiu HW, Chen CH, Chen YJ, Hsu YH. Far-infrared suppresses skin photoaging in ultraviolet B-exposed fibroblasts and hairless mice. PLoS One. 2017;12(3):e0174042. doi:10.1371/journal.pone.0174042

16. Han M, Ban JJ, Bae JS, Shin CY, Lee DH, Chung JH. UV irradiation to mouse skin decreases hippocampal neurogenesis and synaptic protein expression via HPA axis activation. Sci Rep. 2017;7(1):15574. doi:10.1038/s41598-017-15773-z

17. Šimek M, Rubanová D, Nešporová K, et al. Pharmacokinetics of the systemic application of hyaluronic acid for joint arthritis treatment. International Journal of Biological Macromolecules. 2025;307:141937. doi:10.1016/j.ijbiomac.2025.141937

18. Schneider CA, Rasband WS, Eliceiri KW. NIH Image to ImageJ: 25 years of image analysis. Nat Methods. 2012;9(7):671–675. doi:10.1038/nmeth.2089

19. Bankhead P, Loughrey MB, Fernández JA, et al. QuPath: Open source software for digital pathology image analysis. Sci Rep. 2017;7(1):16878. doi:10.1038/s41598-017-17204-5

20. Häder DP, Erzinger GS, eds. Bioassays: Advanced Methods and Applications. Elsevier; 2018.

21. Stirling DR, Swain-Bowden MJ, Lucas AM, Carpenter AE, Cimini BA, Goodman A. CellProfiler 4: improvements in speed, utility and usability. BMC Bioinformatics. 2021;22(1):433. doi:10.1186/s12859-021-04344-9

22. Šimek M, Hermannová M, Šmejkalová D, et al. LC–MS/MS study of in vivo fate of hyaluronan polymeric micelles carrying doxorubicin. Carbohydrate Polymers. 2019;209:181–189. doi:10.1016/j.carbpol.2018.12.104

23. Simek M, Turkova K, Schwarzer M, et al. Molecular weight and gut microbiota determine the bioavailability of orally administered hyaluronic acid. CARBOHYDRATE POLYMERS. 2023;313:120880. doi:10.1016/j.carbpol.2023.120880

24. Šínová R, Pavlík V, Šimek M, et al. The hyaluronan metabolism in the UV -irradiated human epidermis and the relevance of in vitro epidermal models. Experimental Dermatology. 2023;32(10):1694–1705. doi:10.1111/exd.14875

25. Gyongyosi N, Lorincz K, Keszeg A, et al. Photosensitivity of murine skin greatly depends on the genetic background: clinically relevant dose as a new measure to replace minimal erythema dose in mouse studies. Exp Dermatol. 2016;25(7):519–525. doi:10.1111/exd.12984

26. Berton TR, Pavone A, Fischer SM. Ultraviolet-B Irradiation Alters the Cell Cycle Machinery in Murine Epidermis In Vivo. Journal of Investigative Dermatology. 2001;117(5):1171–1178. 10.1046/j.0022-202x.2001.01536.x

27. Hong SP, Kim MJ, Jung M young, et al. Biopositive Effects of Low-Dose UVB on Epidermis: Coordinate Upregulation of Antimicrobial Peptides and Permeability Barrier Reinforcement. Journal of Investigative Dermatology. 2008;128(12):2880–2887. 10.1038/jid.2008.169

28. Inomata S, Takada K, Tsunenaga M, et al. Possible Involvement of Gelatinases in Basement Membrane Damage and Wrinkle Formation in Chronically Ultraviolet B-exposed Hairless Mouse. Journal of Investigative Dermatology. 2003;120(1):128–134. 10.1046/j.1523-1747.2003.12021.x

29. Heckman CJ, Chandler R, Kloss JD, et al. Minimal Erythema Dose (MED) Testing. J Vis Exp. 2013;(75):50175. doi:10.3791/50175

30. Clydesdale GJ, Dandie GW, Muller HK. Ultraviolet light induced injury: Immunological and inflammatory effects. Immunology & Cell Biology. 2001;79(6):547–568. doi:10.1046/j.1440-1711.2001.01047.x

31. Salminen A, Kaarniranta K, Kauppinen A. Photoaging: UV radiation-induced inflammation and immunosuppression accelerate the aging process in the skin. Inflamm Res. 2022;71(7):817–831. doi:10.1007/s00011-022-01598-8

32. Man MQ, Lin TK, Santiago JL, et al. Basis for Enhanced Barrier Function of Pigmented Skin. Journal of Investigative Dermatology. 2014;134(9):2399–2407. doi:10.1038/jid.2014.187

33. Baquié M, Kasraee B. Discrimination between cutaneous pigmentation and erythema: Comparison of the skin colorimeters Dermacatch and Mexameter. Skin Research and Technology. 2014;20(2):218–227. doi:10.1111/srt.12109

34. Matias AR, Ferreira M, Costa P, Neto P. Skin colour, skin redness and melanin biometric measurements: comparison study between Antera® 3D, Mexameter® and Colorimeter®. Skin Research and Technology. 2015;21(3):346–362. doi:10.1111/srt.12199

35. Mah LJ, El-Osta A, Karagiannis TC. γH2AX: a sensitive molecular marker of DNA damage and repair. Leukemia. 2010;24(4):679–686. doi:10.1038/leu.2010.6

36. Wang PW, Hung YC, Lin TY, et al. Comparison of the Biological Impact of UVA and UVB upon the Skin with Functional Proteomics and Immunohistochemistry. Antioxidants. 2019;8(12):569. doi:10.3390/antiox8120569

37. Biernacki M, Skrzydlewska E. Metabolic pathways of eicosanoids—derivatives of arachidonic acid and their significance in skin. Cell Mol Biol Lett. 2025;30(1):7. doi:10.1186/s11658-025-00685-y

38. Dai G, Freudenberger T, Zipper P, et al. Chronic Ultraviolet B Irradiation Causes Loss of Hyaluronic Acid from Mouse Dermis Because of Down-Regulation of Hyaluronic Acid Synthases. The American Journal of Pathology. 2007;171(5):1451–1461. doi:10.2353/ajpath.2007.070136

39. Röck K, Joosse SA, Müller J, et al. Chronic UVB-irradiation actuates perpetuated dermal matrix remodeling in female mice: Protective role of estrogen. Sci Rep. 2016;6(1):30482. doi:10.1038/srep30482

40. Pyun HB, Kim M, Park J, et al. Effects of Collagen Tripeptide Supplement on Photoaging and Epidermal Skin Barrier in UVB-exposed Hairless Mice. Prev Nutr Food Sci. 2012;17(4):245–253. doi:10.3746/pnf.2012.17.4.245

41. Guerrero-Juarez CF, Plikus MV. Emerging nonmetabolic functions of skin fat. Nat Rev Endocrinol. 2018;14(3):163–173. doi:10.1038/nrendo.2017.162

42. Wong Y, Nakamizo S, Tan KJ, Kabashima K. An Update on the Role of Adipose Tissues in Psoriasis. Front Immunol. 2019;10:1507. doi:10.3389/fimmu.2019.01507

43. Kim EJ, Kim YK, Kim MK, et al. UV-induced inhibition of adipokine production in subcutaneous fat aggravates dermal matrix degradation in human skin. Sci Rep. 2016;6(1):25616. doi:10.1038/srep25616

44. Lan CCE, Hung YT, Fang AH, Ching-Shuang W. Effects of irradiance on UVA-induced skin aging. Journal of Dermatological Science. 2019;94(1):220–228. 10.1016/j.jdermsci.2019.03.005

45. Pérez Ferriols A, Aguilera J, Aguilera P, et al. Determination of Minimal Erythema Dose and Anomalous Reactions to UVA Radiation by Skin Phototype. Actas Dermosifiliogr. 2014;105(8):780–788. doi:10.1016/j.adengl.2014.05.020

