## Supplementary figures for "Characterization of a chronic UV-induced photoaging mouse model: insights into skin barrier dysfunction, extracellular matrix remodeling, and altered adipogenesis"

### S. Fig. 1

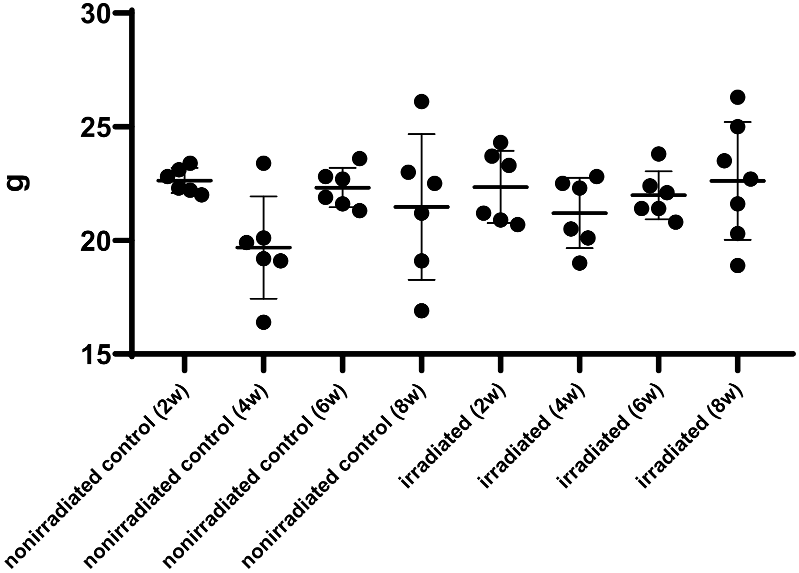


**Supplementary Figure 1: Weight of the mice.** At the beginning of the experiment, all mice were weighed and randomly assigned to eight experimental groups - nonirradiated controls sacrificed at 2 (N=6), 4 (N=6), 6 (N=6), and 8 (N=5) weeks and irradiated mice sacrificed at 2 (N=6), 4 (N=6), 6 (N=6), and 8 (N=7) weeks post-exposure.
